# Reverse development in the ctenophore *Mnemiopsis leidyi*

**DOI:** 10.1101/2024.08.09.606968

**Authors:** Joan J. Soto-Angel, Pawel Burkhardt

## Abstract

Reverse development, or the ability to rejuvenate by morphological reorganization into the preceding life cycle stage is thought to be restricted to a few species within Cnidaria. To date, the cnidarian *Turritopsis dohrnii* is the only known species capable of undergoing reverse development after the onset of sexual reproduction. Here, we demonstrate that the ctenophore *Mnemiopsis leidyi* is capable of reversal from mature lobate to early cydippid when fed following a period of stress. Our findings illuminate central aspects of ctenophore development, ecology, and evolution, and show the high potential of *M. leidyi* as a new model system to study reverse development and rejuvenation. Besides shedding light on the plasticity of developmental programs, our results raise fundamental questions about early animal development, body plans and life cycles.

## Introduction

Marine invertebrates display an outstanding diversity of life cycle strategies which greatly differ not only between main groups, but also among closely-related species (1). Addressing how ecology and evolution interplay and shape life histories remains a great challenge. The scarce knowledge on the full repertoire of life cycles for most marine invertebrates obscures how phylogenetically spread certain traits are, which in turn hinders a profound understanding of central aspects of animal development, ecology and evolution. To date, the capacity of one life cycle stage to transform back to the preceding stage by morphological reorganization has been regarded as a distinctive and unparallelled feature of cnidarians (2). This ability for reverse development known for a few cnidarian species was first reported over a century ago, and gained wide renown with the discovery of the peculiar life cycle of the so-called immortal jellyfish, *Turritopsis dohrnii* (2-6). This hydrozoan is currently considered the only animal able to repeatedly rejuvenate after sexual reproduction, challenging our understanding of aging and suggesting a potential for biological immortality (6).

Ctenophores (or comb jellies) are one of the oldest extant animal lineages. Accumulated evidence supporting their phylogenetic position as the sister group to all other animals (7) place them as a pivotal model to study unique evolutionary innovations potentially rooted within the deepest branches of the animal tree of life (8). Their extreme fragility stands as one of the main reasons behind the limited available knowledge on their true diversity and biology. Ctenophore life cycle strategies and their potential plasticity are still insufficiently explored to draw reliable conclusions at phylum level (9). The typical ctenophore life cycle includes a planktonic, bitentaculate cydippid stage, also present in benthic species within Platyctenida, but absent in Beroida. Most lobate ctenophores hatch as cydippid larva and undergo a gradual metamorphosis in which the tentacles are reabsorbed, and newly-developed paired structures appear (i.e. auricles and lobes) (10, 11). Prior studies with one of the best-known ctenophore species, *Mnemiopsis leidyi*, have shown that starvation alone is insufficient to initiate reverse metamorphosis from lobate to cydippid (11). To date, reverse development has not been documented in ctenophores. Here, using individual trajectories of morphometric changes over time for physiologically stressed animals, we provide the first evidence of reverse development in Ctenophora.

## Results

To determine if ctenophores can undergo reverse development, we assessed two different stressors; prolonged starvation and physical injury (lobectomy), both followed by a low feeding regime (**Figure 1A, B**) (for details see Materials & Methods section). Developmental changes of main anatomical features were monitored over time (**Figure 1B-F**). Animals were considered fully reversed when they exhibited a typical cydippid morphology, i.e., rounded body, absence of lobes and auricles, and presence of two fully formed, functional tentacles (**Figure 2, Supplementary Video 1**).

**Figure 1.**
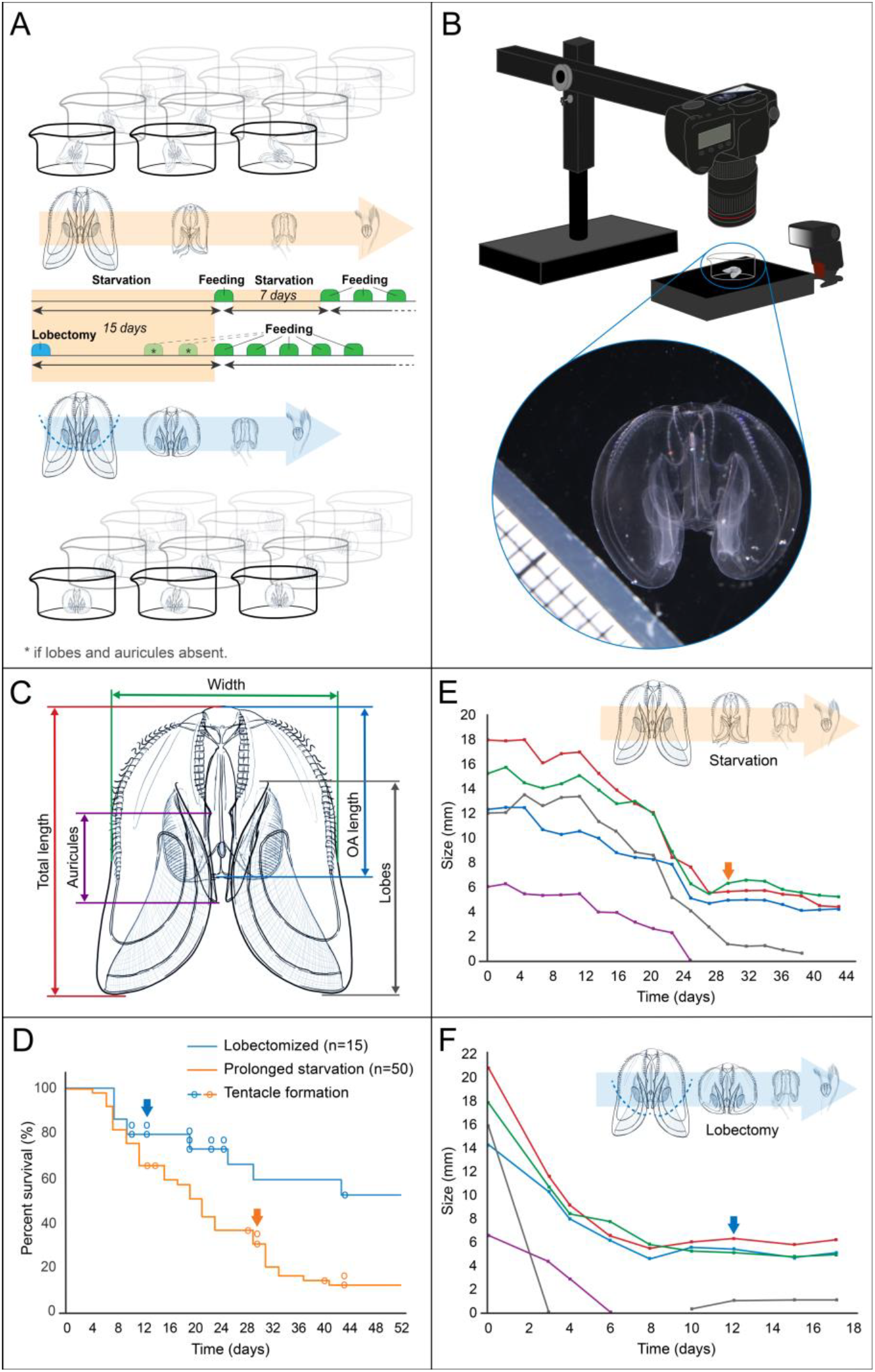
Reverse development in the ctenophore *Mnemiopsis leidyi*. (A) Experimental design for starved (above) and lobectomized (below) *M. leidyi*. (B) Imaging station to record morphometric changes over time. (C) Detail and extent of the anatomical features monitored over the course of the experiments. (D) Percentage of survival over time across the two treatments. Circles over the line indicate tentacle regeneration for all individuals that displayed reverse development (n=20). Arrows correspond to the animals shown in panels E and F. (E) Morphometric changes for a starved individual. (F) Morphometric changes for a lobectomized individual. Arrows point to tentacle regeneration. Colour code for anatomical features followed in panels E and F as shown in panel C.

**Figure 2.**
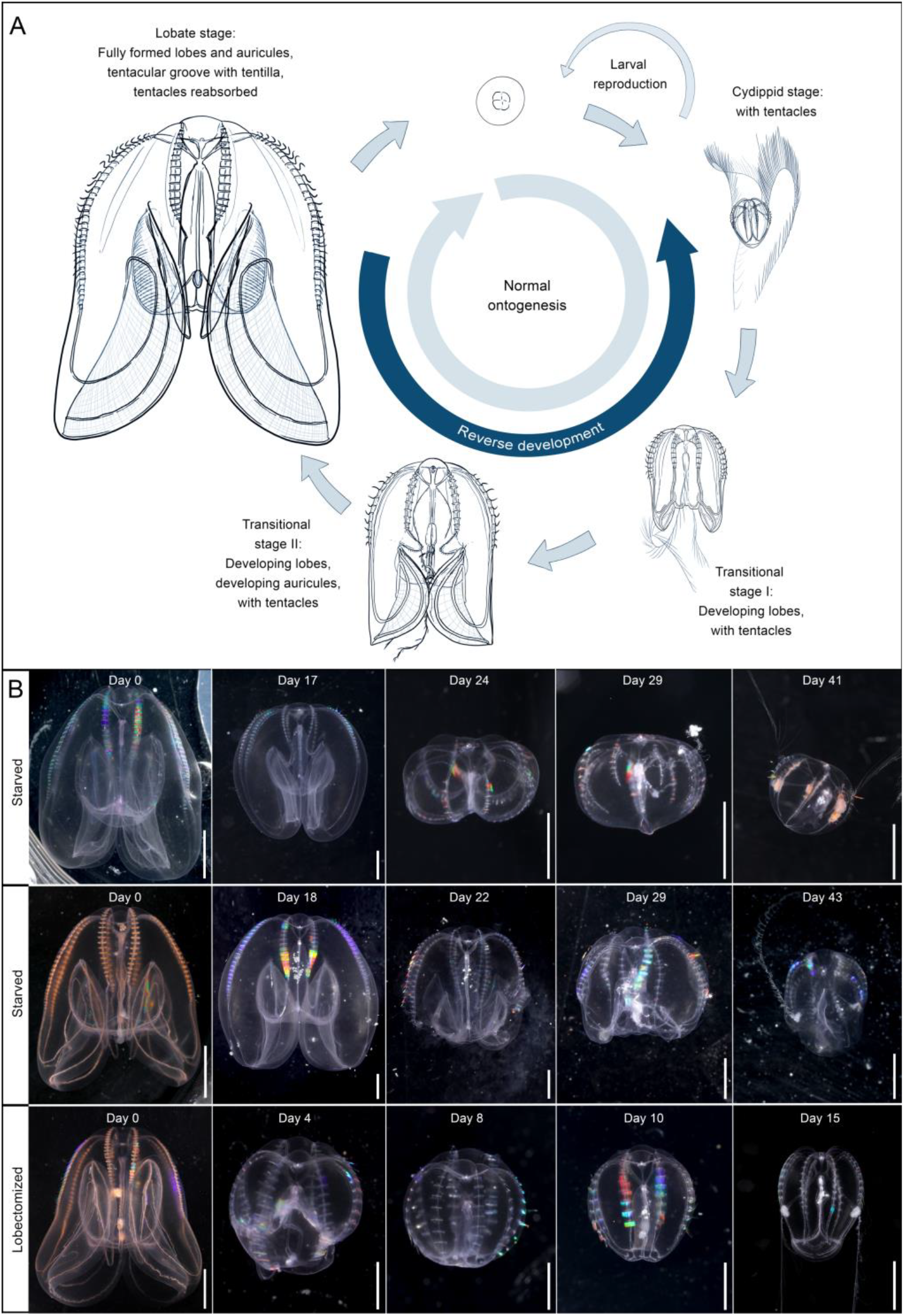
Life cycle and main morphological changes of the ctenophore *Mnemiopsis leidyi*. (A) Ordinary downstream development (normal ontogenesis, clockwise) and reverse development (anticlockwise). Note the absence of tentacles in the fully transitioned lobate stage, and the presence of newly developed anatomical features (i.e. auricles and lobes) gradually appearing during metamorphosis of the cydippid stage and shrinking until disappearing during reverse development. Illustrations of the different life cycle stages by Nicholas Bezio. (B) Individual trajectories and morphological changes during reverse development for three *M. leidyi* specimens (two starved and one lobectomized) that fully reversed to a typical bitentaculate cydippid stage. Note the increase of prey items in the gut when tentacles regenerated. Scale bar: 5 mm for Day 0; all others 2 mm.

Both treatments were found to trigger reverse development, with 13 animals out of 65 used showing complete lobate-cydippid reversion. Seven additional animals developed tentacles but only partially reabsorbed the lobes and/or auricles. We noted that lobectomized animals shrank and reversed faster than only starved, with tentacles regained on day 20 ± 9 at 5.3 ± 1.0 mm (OA length, n=12) vs 30 ± 12 days at 3.5 ± 1.3 mm (OA length, n=8), respectively (**Figure 1D**).

Interestingly, lobectomy treatment resulted in lower mortality (**Figure 1**) and higher reversion success, with six out of fifteen (40%) animals successfully reversed (80% when including partially reversed animals). In contrast, prolonged starvation only yields into seven out of 50 (14%) fully reversed animals (16% when considering partially reversed).

In most cases, the auricles regressed faster than the lobes (**Figure 1E, F**), and were completely resorbed while the lobes were still present. In some instances, the tentacular bulbs and the mouth area were completely lost after partial disintegration, but were subsequently regenerated a few days after being fed (**Figure 2**). Fully reversed animals showed typical cydippid behavior, being able to efficiently capture, manipulate and ingest *Brachionus* (Rotifera) using their tentacles as described in (12). Later, they further developed to normal lobate *M. leidyi* able to produce viable gametes. We noted that fully reverted individuals (12 out of 13) also spawned while having a cydippid morphology, as expected for animals in that size range (13, 14). Overall, 20 animals (31%) developed fully formed, functional tentacles *de novo* after being lost during normal ontogenesis.

## Discussion

Conventionally, cnidarians have been considered the sole animal group capable to undergo reverse development (2, 4), with only a few species reported to date. Examples of non-metagenetic cnidarians (i.e. species with no alternance of sexual and asexual generations) are restricted to one known case of coral in which the primary polyp is known to potentially transform back into a free-swimming, planktonic, planula larva (2, 4). Metagenetic cnidarians (i.e. species with both medusa and polyp stage) with known reverse development capabilities are restricted to a few species in which a recently produced medusa transforms back into a polyp (2, 4). So far, *T. dohrnii* is the only species in which fully-grown, sexually mature medusae can revert into a polyp stage (3). Besides these cnidarian species, an additional case have been reported as a putative case of reverse development within Platyhelmintha: the parasitic cestode *Echinococcus granulosus* (15). Protoscoleces stages of *E. granulosus* preceding the scolex stage that in turn develop into the adult stage are capable to encyst secondarily after escaping from a ruptured hydatid cyst (15, 16). Our finding of ontogeny reversal in Ctenophora confirms that reverse development might be more widespread than previously thought. Admittedly, reverse development in Cnidaria and Ctenophora are markedly different in several aspects. In Cnidaria, the transition occurs between discrete life stages (medusa to polyp, or polyp to planula) and by means of a profound tissue reorganization, resulting in the generation of a new organism from a pool of poorly-differentiated cells (3). In contrast, reverse development in *M. leidyi* occurs gradually, within the same individual, and between life stages showing a continuum: lobate, transitional and cydippid stages (**Figure 2**). Despite these differences, the process described here fulfills the definition of reverse development as originally described for Cnidaria: a shift in the ordinary, downstream development into the opposite ontogenetic direction by back-transformation of some life stages (2). While the existence of a true larval stage in Ctenophora is under debate (9, 13), the normal ontogeny of lobate ctenophores unquestionably proceeds from the early cydippid to later lobate stage. Metagenetic species with alternance of generations constitute a great majority of the documented cases of reverse development. Our findings using *M. leidyi* represent the first case of a non-metagenetic species in which the same reproductive individual is capable of gradually rejuvenate to an earlier immature stage.

Our results suggest that reverse development in *M. leidyi* can be triggered by at least two stressors: prolonged starvation and injury. Similarly, the jellyfish *T. dohrnii* undergoes reverse development in response to injury, adverse environmental conditions, or aging (17). It is therefore possible that some early-stage individuals of *M. leidyi* observed in the field were previously fully-formed adults, thus challenging the current knowledge on the population dynamics of the species. The ability *of M. leidyi* to considerably reduce size and body mass through starvation was already known (11, 18). However, previous starvation experiments concluded that animals cannot reverse metamorphosis to go back to the cydippid larval stage (11). Therefore, it seems that starvation alone is not enough to induce reverse development. Our results also support this, as most of the animals starved for a prolonged period were too weak to recover and perished, while the ones that successfully reversed to cydippid stage only developed tentacles after being fed. The high body plasticity and tolerance for starvation in *M. leidyi* has been pinpointed as a potential mechanism to overcome periods of low food availability, and a key reason for its success as a highly invasive species (11, 18). Reverse development not only entails a reduction in size, but a change in morphology and associated feeding mode. The diet of wild animals dramatically changes over the course of normal development, from a predominantly microplankton diet (protists) to a metazoan-based diet (19, 20). Therefore, when regressed animals regain feeding structures (tentacles) that were lost due to ontogenetic changes, they gain access to food resources *de novo* that were not available to or less efficiently captured by the lobed adults (10, 12, 19), with a subsequent ecological niche reversal. This finding adds a new perspective on a wide range of ecological aspects, such as those evaluating the high dispersal capabilities and invasive success in *M. leidyi* (11).

The occurrence of reverse development in *M. leidyi* poses fundamental questions on the molecular and cellular mechanisms involved. In *T. dohrnii*, this is mediated by cell trans-differentiation (3), with high activity and/or expansion of genes involved in aging/lifespan, the regulation of transposable elements, DNA repair, telomere maintenance, redox environment, stem cell population and chromatin remodeling (5, 6). While fundamental knowledge on the cellular and molecular mechanisms underlying the extreme regeneration capabilities of *M. leidyi* exist, specific pluripotent stem cells have not yet been identified (21-23). Ctenophore tentacular bulbs are the main regions of cell proliferation, but are not required for regeneration (21). To which extent reverse development and regeneration share common attributes on cell reprogramming and molecular pathways remains an open question. This study opens new avenues for research using *Mnemiopsis leidyi* as a model species to study aging and rejuvenation in animals.

The discovery of an alternative, upstream development that ultimately leads to escape the normal ontogenetic fate (i.e. senescence) in *M. leidyi* has broad implications on the plasticity of developmental programs in early-diverging animals (13). In this sense, reverse development in a ctenophore raises fascinating questions about its phylogenetic extent, evolutionary origin and the evolution of life histories within and beyond Ctenophora.

## Materials and Methods

### Ctenophore culture

All animals used were healthy, 2-3 months old, reproductive *M. leidyi* specimens raised as described in (24). Before the start of the experiment, specimens selected were carefully inspected under the microscope and only fully transitioned individuals, ca. 2 cm in total length or larger, with completely resorbed tentacles were selected (11, 19). Animals were kept individually in 300 mL crystallizing dishes provided with a lid to avoid evaporation. Filtered seawater as described in (24) was used and changed every 1-3 days, allowing water parameters (temperature 20ºC ± 0.5; salinity 27 ± 0.5 ppt; pH 8.0 ± 0.1) to remain virtually constant during the entire timeframe of the experiments.

### Experimental design

Two treatments simulating different stress conditions were assayed: prolonged starvation (n=50) and physical damage (n=15). For the latter, damage was inflicted by surgically removing the lobes at the level of the mouth. For both treatments, the animals were initially starved for 15 days, and later fed once a week with a small amount of *Brachionus* (Rotifera), ca. 100-200 prey items per crystallizing dish, equivalent to ca. 0.5-1 prey/ml. For details on *Brachionus* culture and feeding see (24). Ctenophores were fed every second day once the lobes were fully reabsorbed. For lobectomized animals shrinking quickly, this translated into feeding before the end of the initial 15 days period of starvation. The most critical point arises when specimens have no lobes or tentacles, as they become extremely delicate. At this stage animals must be fed, but they show very low feeding success and rarely catch prey. Occasionally, animals without tentacles or lobes are still able to passively consume rotifers that accidentally get trapped by the mucous secreted in the mouth area. This is enough for them to regenerate the tentacles (and tentacular bulbs when missing). When tentacles are regenerated, the number of *Brachionus* observed in the gut increases considerably. This can be observed in Figure 2B.

### Data acquisition

Morphometric changes were recorded every 1-3 days for both experiments. In addition, appearance/reabsorption of morphological structures, partial disintegration, sudden damage, or occurrence of sexual reproduction (i.e. egg production) were monitored. For specificities on the morphological features monitored see Figure 1C. When paired structures (i.e. lobes and auricles) were visibly asymmetrical, the longest one was measured. Animals were carefully transferred individually to 50 mL crystalizing dishes as described in (24) and photographed against a black background and a millimetric scale. Photography gear consisted of a Canon EOS 5D Mark IV, Canon MP-E65mm f/2.8 1-5x Macro lens and Canon Speedlite 430EX II flash. Relative position of the different gear elements as shown in Figure 1B. Once photographed, ctenophores were placed back in a clean 300 ml crystallizing dish with renovated seawater. Measurements were obtained through Adobe Photoshop 2023.

## Supporting information

Timelapse showing reverse development in a lobectomized individual of M. leidyi.

## Acknowledgments

We thank Nicholas Bezio for the illustrations of *M. leidyi* life cycle, and Jeffrey Colgren and Aino Hosia for their valuable comments on the manuscript. This work was supported by the Michael Sars Centre core budget and the European Research Council Consolidator Grant (101044989, “ORIGINEURO”) awarded to P.B.

## Author Contributions

JJSA and PB designed research; JJSA performed research; JJSA and PB contributed new reagents/analytic tools; JJSA and PB analyzed data; and JJSA and PB wrote the paper.

## Supplementary files

**Supplementary Video 1**: Timelapse showing reverse development in a lobectomized individual of *M. leidyi*.

